# Counting to two: how phages decide between lysis and lysogeny

**DOI:** 10.64898/2026.05.14.725151

**Authors:** Janni Harju, Ghita Guessous, Zemer Gitai, Ned S. Wingreen

## Abstract

Upon infecting a bacterium, temperate phages must decide between killing the cell to reproduce (lysis) and entering a symbiotic lifestyle (lysogeny). This choice is often informed by the cells state and by the number of infecting phage particles, or multiplicity of infection (MOI). Because the copy numbers of all phage genes scale identically with MOI, an MOI-dependent decision requires a rapidly established asymmetry between the lytic and lysogenic pathways. We introduce a minimal model showing how a simple coupling to host machinery (e.g. a protease) can generate such an asymmetry. Our analytical model yields scaling laws that predict how the lysislysogeny decision responds to perturbations of the host. It also quantitatively predicts stochastic features of infection outcomes, such as lysogeny probabilities, consistent with experimental data. By reducing complex regulatory networks to their essential components, our framework reveals the organizing principles of lysislysogeny decisions across phage species.

## I. INTRODUCTION

Temperate phages are bacterial viruses that must decide between two alternate life-paths upon entering a host. This decision is generally informed by the number of phage particles infecting the host: if the multiplicity of infection (MOI) is one, the phage typically replicates and then lyses the bacterium, releasing a burst of progeny. However, at higher MOIs, which indicate a higher ratio of phage particles to susceptible hosts in the local environment, the phage can instead integrate its genome into the bacterial cell, forming a lysogen [1].

The lysis-lysogeny decision mechanism of the *Escherichia coli* phage *λ* has been extensively studied for over six decades [2, 3]. Early works based on mutant assays identified key genes required for each infection pathway. High levels of the *λ* protein CII promote both integration and expression of CI, a repressor that maintains the lysogenic state. At low MOIs, this pathway is suppressed by degradation of CII by the essential host protease FtsH [4, 5].

Interestingly, more recent studies have identified novel lysis-lysogeny decision mechanisms that are also coupled to host FtsH activity [6, 7]. Both these and the *λ* decision pathway have been successfully modeled *in silico* [7–12]. While these models can reproduce the MOI dependence of post-infection dynamics, their complexity makes it difficult to extract any general principles that may underpin lysis-lysogeny decisions across species. Furthermore, questions remain about how MOI-independent early gene expression dynamics can give rise to MOI-dependent lysis-lysogeny outcomes [12].

A complete model for temperate phage decisions should also explain why infection outcomes vary significantly at the single-cell level [13–18]. Such variations are thought to partially result from variations in host state, for instance, cell size [14, 15]. However, stochastic fluctuations [10, 19] and differences in infection timing [11, 12] are also thought to play a role. The effects of these sources of noise have been modeled in *λ* [10, 11, 19], but it remains unclear to what extent these models and their conclusions might generalize across phage species. Here, we introduce a general coarse-grained model that lays out minimal requirements for an MOI-dependent lysis-lysogeny switch. The model requires a mechanism that controls protein accumulation dynamics on relevant, short timescales and in a targeted manner. Motivated by the established role of FtsH in known systems, we consider protease-mediated degradation of lysogeny-inducing proteins as an illustrative example mechanism. Our generalized framework predicts scaling laws describing how lysis-lysogeny decisions shift with host state. Additionally, we introduce an analytical approach for understanding the stochastic nature of infection outcomes, as well as the effects of infection timings. Our work suggests that phage decision making mechanisms may share universal features and helps identify the most consequential interactions in such decision pathways.

## II. RESULTS

### A. Scaling arguments for lysis-lysogeny mechanisms

To construct our model, we assume that the lysis-lysogeny decision is based on the relative dynamics of two competing pathways (Fig. 1a). At time *t* = 0, the concentration of phage genomes *g*_phage_ inside the bacterial host is proportional to the MOI. The phage genome encodes a lysis-promoting and a lysogeny-promoting protein, whose corresponding mRNA and protein concentrations are denoted by *m*_lyt_*/m*_lyso_ and *p*_lyt_*/p*_lyso_. We assume that each protein has a designated decision threshold concentration 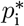 [10, 12]. If a protein reaches this level, it activates its own pathway and represses the alternate one. The simplest dynamics for these components consist of transcription, translation, as well as protein and mRNA degradation and dilution (see Appendix A). However, since the lysis-lysogeny decision of *λ* occurs within the first *∼*5 min post-infection, not all of these processes act on relevant timescales. Degradation rates for *λ* transcripts are estimated at 1*/*10 min^*−*1^ [12]. Protein dilution and degradation rates in bacteria are generally on the order of 1*/*20 min^*−*1^ [20]. Hence, on timescales *t ≲*5 min, the protein and mRNA dynamics are well approximated by

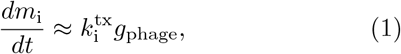

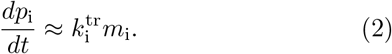

**FIG. 1.**
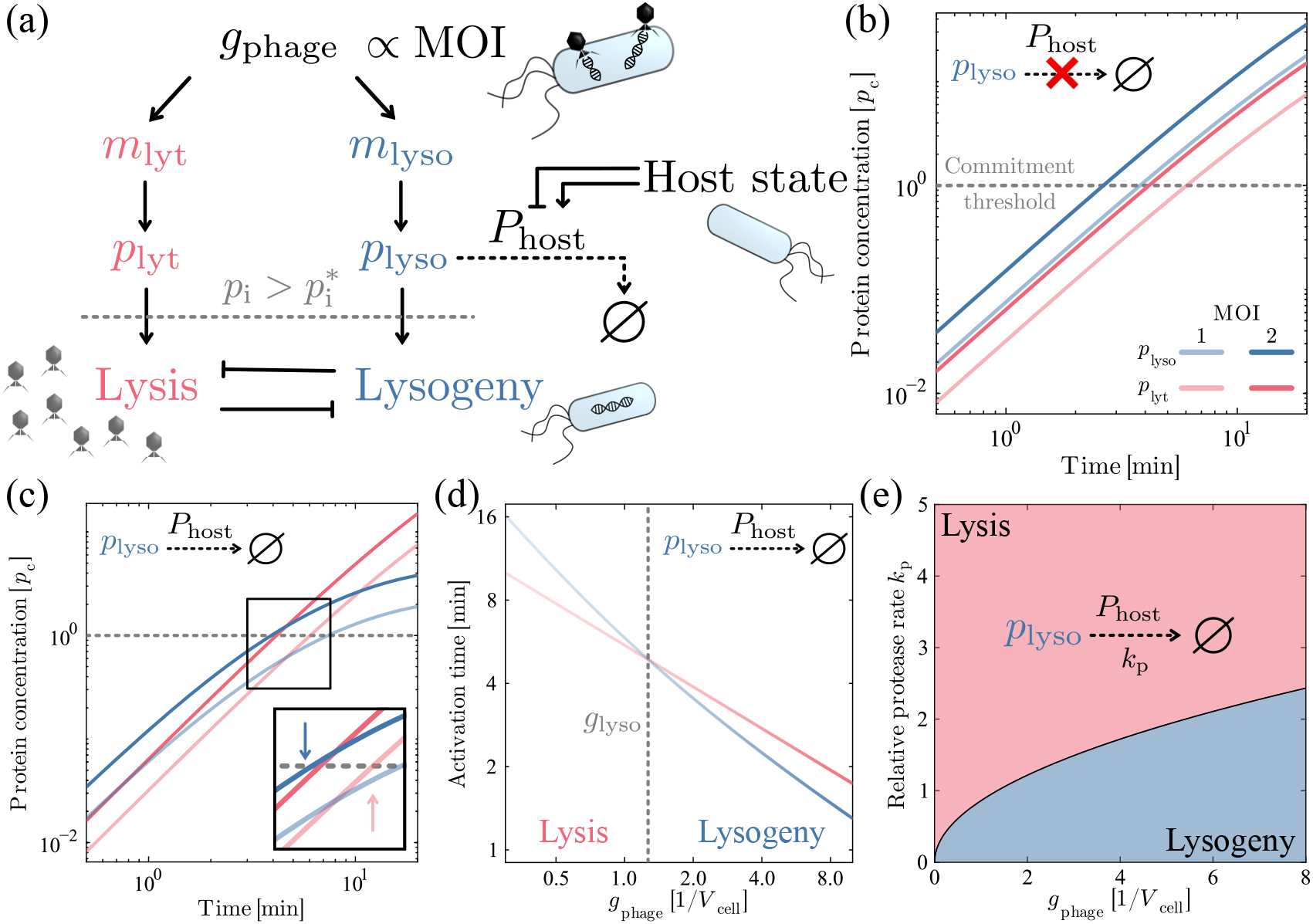
Minimal model for lysis-lysogeny decisions. (a) Illustration of the model. mRNAs for both a lysisand a lysogeny-promoting protein are transcribed at rates proportional to the multiplicity of infection (MOI). Once either protein concentration, *p*_lyt_ or *p*_lyso_, reaches a commitment threshold, it triggers its own pathway as well as repression of the alternative pathway. An asymmetry is introduced by host protease-mediated degradation of the lysogeny-promoting protein. (b) Protein dynamics in the absence of host protease-mediated degradation; on short timescales, both protein levels increase quadratically in time, for all MOIs. Parameters were set following [12] (Table I). mRNA degradation decreases temporal exponent on timescales 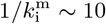 min. (c) Protein dynamics with host protease-mediated degradation of the lysogeny-promoting protein at rate *k*_p_. At timescales *t ∼* 1*/k*_p_, *p*_lyso_ transitions to a linear scaling with time. Inset shows that the two protein curves now cross, and a change in the MOI can change which protein reaches the threshold level first. (d) Scalings of the times to reach commitment threshold with respect to the phage genome concentration *g*_phage_ within the cell. Above *g*_lyso_, the outcome is lysogeny. At large *g*_phage_, the threshold is reached at *t <* 1*/k*_p_. Concentration units are in phage genomes per cell volume *V*_cell_. (e) Phase diagram of the model. Black curve represents *g*_lyso_(*k*_p_). In the lytic regime, degradation of *p*_lyso_ is fast enough compared to transcription and subsequent translation to prevent lysogeny. Unless otherwise stated, all results are shown including mRNA degradation at rate 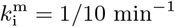. Results are similar with 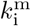 (Supplemental Figure S1a,b).

**TABLE I.**
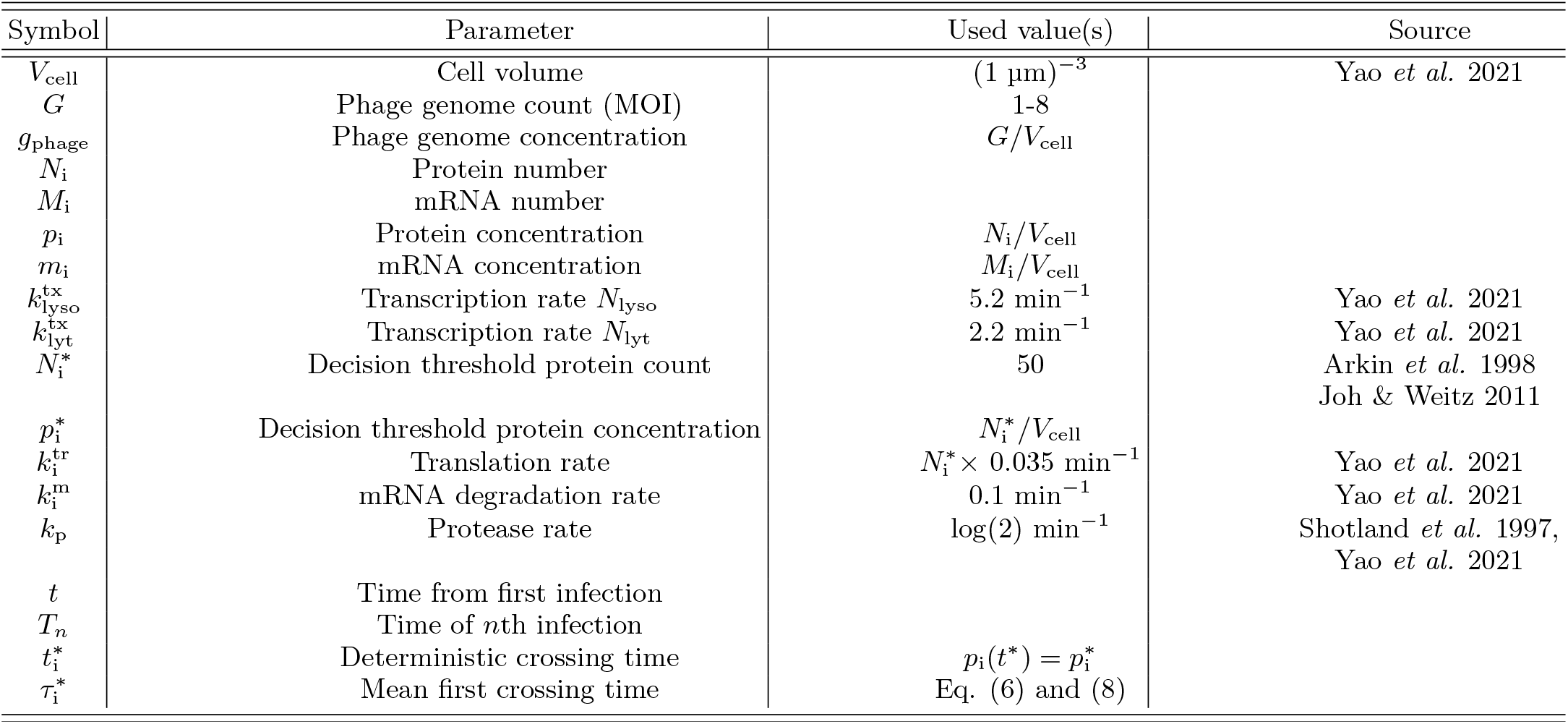
List of all symbols and parameter values used.

Here, i is the index lyt or lyso, and 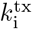 and 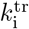 are the transcription and translation rates, respectively. mRNA concentrations therefore increase approximately linearly in time 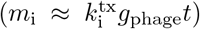, and protein levels increase quadratically 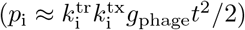.

To visualize the effects of MOI changes on phage protein dynamics, we plot *p*_i_(*t*) on logarithmic axes, where the slope of each curve is set by the exponent of *t*; here 2 (Fig. 1b). The vertical separation between the two curves is set by the ratio of products of transcription and translation rates of the two mRNA species. Upon a doubling of the MOI, total transcription rates double, and both curves shift upwards by equal amounts. The order of the parallel curves does not change with MOI, and hence the same protein reaches the decision threshold first.

An MOI-dependent decision mechanism thus requires an asymmetry beyond different transcription and translation rates. This additional mechanism should specifically target one pathway on timescales shorter than 5 minutes. One possibility would be a nonlinearity in Eqs. (1) or (2), corresponding to feedback between mRNA/protein levels and transcription/translation rates. Alternatively, either pathway might couple to host- or phage-encoded machinery. Any phage-encoded machinery would first have to be transcribed and translated. Since slower post-infection dynamics would give the host more time to mount an immune response, coupling to host-encoded machinery is expected to be favored. Moreover, by basing decisions on host-encoded factors, phage can potentially gain information about their host’s state (see Discussion).

As a biologically motivated example, we consider degradation of the lysogeny-inducing protein by a host protease, e.g. FtsH in *E. coli*. Bacterial proteases are highly specific, and can degrade proteins on timescales of *∼* 2 min [5]. We hence introduce a protease degradation term:

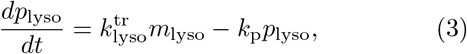

where *k*_p_ is the protease degradation rate. At times *t ≲*1*/k*_p_, *p*_lyso_ still increases quadratically. However, for *t »* 1*/k*_p_, the scaling of *p*_lyso_ becomes linear (Fig. 1c). This implies that the concentration curves *p*_i_(*t*) can now intersect. Critically, an increase in the MOI, e.g. from 1 to 2, can now move the crossing point of the two curves across the 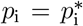 line. As a result, a different protein – lytic or lysogenic – can reach the decision threshold first (Fig. 1C inset) while the other protein’s concentration is still *∼* 20% below its decision threshold (see Appendix B). Intuitively, at larger MOIs, transcription of *m*_lyso_ is sufficiently fast to outpace protease-mediated degradation, thus leading to lysogeny.

Within our model, the decision switches at a genome concentration where *p* _lyt_ and *p* _lyso_ reach their target levels simultaneously (Fig. 1d). Since the crossing point occurs at time *t* _x_*∼*1*/k* _p_ (see Appendix C), setting 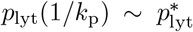, we find a scaling prediction for the minimum phage genome concentration for lysogeny:

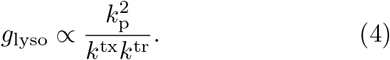

Here, *k*^tx^ and *k*^tr^ are the baseline transcription and translation rates of the phage genes, and a constant prefactor is set by the ratios of 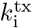, 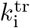, and 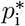 (see Appendix D).

Eq. (4) shows that a 30% decrease in the protease degradation rate could shift the MOI at which lysogeny occurs from two to one, effectively preventing lysis (Fig. 1e).

### B. Model predicts decision stochasticity

Our scaling argument was derived on the population average level, neglecting stochastic effects. On average, for *g < g*_lyso_, we expect lysis, and for *g > g*_lyso_, lysogeny.

However, experiments have revealed substantial stochasticity of outcomes on a single-cell level, with approximately 20-40% of cells infected by a single *λ* phage becoming lysogens [15]. Here, we show that our minimal model predicts similar levels of outcome variability.

To study the effects of stochasticity, we conduct Gillespie simulations of our model with rates consistent with early-stage *λ* infections [12] (Appendix E). These simulations can be used to sample noisy mRNA and protein trajectories at the single-cell level (Fig. 2a) Even in the presence of stochasticity, our formula for *g*_lyso_ (Eq. (4)) accurately predicts the preferred outcome at different MOIs and protease activity rates (Fig. 2b).

**FIG. 2.**
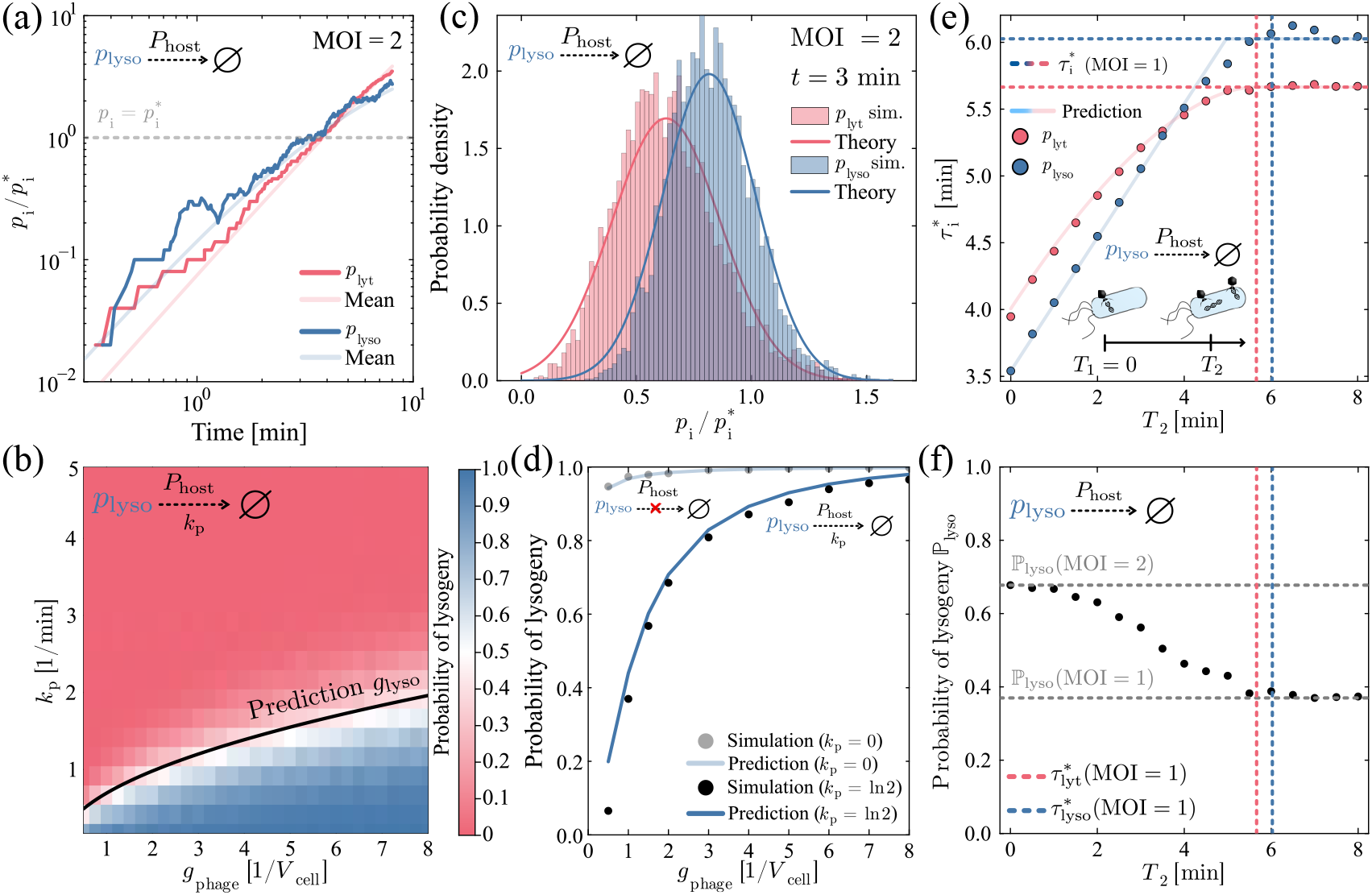
Effects of stochasticity and timing of second infection. (a) An example of a stochastic protein trajectory from a Gillespie simulation with MOI = 2 resulting in lysogeny, overlaid on the mean curves for each protein species. (b) The probability of lysogeny from stochastic Gillespie simulations, at varying MOIs and protease degradation rates. Black curve shows theoretical prediction for *g*_lyso_ (Eq. (4)), the phage genome concentration where the deterministic decision is expected to switch. Results are similar without mRNA degradation (Supplemental Figure S1c). (c) Histograms showing the distribution of single-cell protein concentrations in Gillespie simulations, at MOI = 2, 3 minutes after infection. Overlaid are predictions assuming approximately Gaussian distributions, with the theoretically predicted means and variances. Histograms for mRNA and protein distributions at other time-points are shown in Supplemental Figure S2. (d) Probability of lysogeny from Gillespie simulations, together with theoretical predictions given by Eq. (5). Data are shown in the presence of protease activity, with protein lifetime of 1 minute (dark curve and points) as well as in the absence of protease activity (light curve and points). Results are similar without mRNA degradation (Supplemental Figure S1d). (e) Mean first passage time for each protein as a function of the time *T*_2_ at which a second phage infects the cell. Dashed vertical and horizontal lines indicate the mean first passage times at MOI = 1. Light curves show predictions given by Eq. (11) and (12). Prediction for 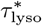 becomes inaccurate when 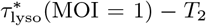 is smaller than 1*/k*_p_, i.e. when second order effects are relevant. (f) Probability of lysogeny as a function of the time at which a second phage infects the cell. Dashed vertical lines again indicate mean first passage times at MOI = 1. Dashed horizontal lines show the probability of lysogeny at MOIs of 1 and 2.

To better understand the stochasticity of infection outcomes, we consider the distribution of protein concentrations as a function of time. In contrast with previous works [21–25], we focus on short timescales postinfection, when neither the mRNA nor protein concentration has equilibrated. For both *p*_lyt_ and *p*_lyso_, analytical expressions for the time-dependent concentration distributions can be found (Appendix F, G). On relevant timescales, we find that the joint (*m*_i_(*t*), *p*_i_(*t*)) distributions are approximately Gaussian (Fig. 2c, Supplemental Figure S2), where the time-dependent means *µ*_i_(*t*) and standard deviations *σ*_*i*_(*t*) can be found exactly (Appendix F, G). At any time *t* when both distributions have some weight above 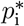, *either* decision may have already been reached. Our simulations show that at MOI = 1, the probability of lysogeny is approximately 40%, and that even at high MOIs, the probability remains below 100% (Fig. 2d). These values and the shape of the curve are similar to experimental data for *λ* [15], without any parameter fitting.

### C. First passage times set lysogenization probability

To develop a theoretical understanding of outcome stochasticity, we note that the probability of lysogeny is set by the statistics of the first passage times at which the lysis and lysogeny proteins reach their decision thresholds [10]. When both proteins accumulate independently before commitment to a decision, the probability of a lysogeny decision is

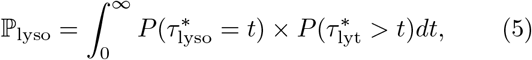

where 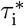 is the first passage time at which protein i crosses its threshold 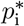 . We integrate over time to find the probability that *p*_lyso_ reaches its decision threshold at time *t* while *p*_lyt_ is still below its decision threshold.

Eq. (5) requires expressions for the first passage time distributions of both protein species. First, we consider *p*_lyt_. Without protein degradation, the probability of a first passage time smaller than *t* can be written as the probability that the concentration at time *t* is above 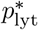:

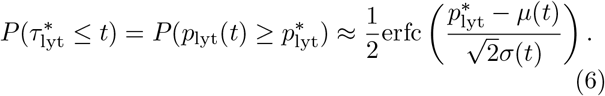

Here erfc is the complementary error function. Differentiating Eq. (6) with respect to time gives the probability density function for 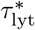 (Appendix H).

Due to protein degradation events, the *p*_lyso_ concentration can cross its decision threshold multiple times. Hence, Eq. (6) does not hold; even if the current concentration is below the decision threshold, the threshold may have been crossed previously. At any given point in time, the expected rate of *p*_lyso_ crossing its threshold is given by the expected translation rate times the probability density at the threshold (Appendix K):

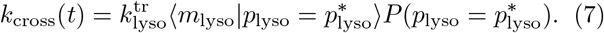

In the limit of approximately Gaussian distributions, the conditional average of the mRNA concentration can be found exactly (Appendix K).

Integrating *k*_cross_ over a time interval yields a mean number of crossings. However, multiple crossing events are expected to occur around the time 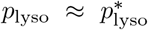 . Hence, to estimate the probability of a set of consecutive crossings occurring at a given time, we use a clump rate approximation [26]. The mean total rate of crossings is approximately the rate of a “clumped” crossing times the number of crossings in a clump, *c*(*t*) (Supplemental Figure S3). We estimate *c* (*t*) by assuming a geometric distribution for escapes from the threshold (Appendix K). Then, the probability of a clump of crossings before time *t* can be found as

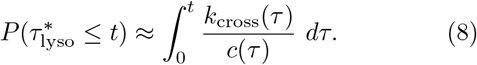

By combining these relations into Eq. (5), we find a theoretical prediction for the probability of lysogeny at a given MOI, consistent with our exact simulation results (Fig. 2d).

### D. Transition between noise regimes governs ℙ _lyso_

Earlier, we noted that in the absence of a mechanism that breaks the symmetry between protein species, we expect the outcome to be independent of MOI (Fig. 1b). However, in the presence of stochasticity, a small percentage of infections can still give rise to the unpreferred outcome even without a symmetry-breaking mechanism, e.g. in the absence of protease activity (Fig. 2d, faint curve). In this case, the probability of lysogeny has a weak MOI dependence, which can be understood as the transition between two noise regimes (Appendix J). At high MOIs, the noise levels saturate to those set by the protein count at the threshold, independent of *g*_phage_. At low MOIs, noise is dominated by mRNA copy number variations, which depend on *g*_phage_, and can thus give rise to a weak MOI-dependence of ℙ_lyso_.

With protease activity, at a genome concentration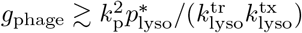 reaches its critical level before protease activity becomes relevant. At high MOIs, the probability of lysogeny thus plateaus at the same levels as without protease activity (Fig. 2d), set by protein count noise. At low MOIs, the first passage times 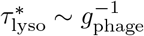 and 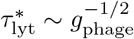 show a cross-over, as discussed earlier, and ℙ_lyso_ drops below 50%. However, the decision remains stochastic due to variations in protein and mRNA copy numbers.

### E. Timing of second infection

So far, we have considered nearly simultaneous infections by multiple phages, corresponding to a constant *g*_phage_ during the decision-making process. We now relax this assumption to derive general predictions for the effects of delayed later infections. Let *T*_*n*_ be the time of the *n*th infection; we set *T*_1_ = 0. At each infection, the phage genome concentration increases by *g*_1_, the phage genome concentration corresponding to an MOI of 1.

Suppose that the mean of *p*_i_(*t*) is order *α* in time; 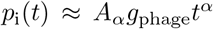, where *A*_*α*_ is some constant. Each phage genome can be seen as an independent source of mRNA and protein, so that the mean protein dynamics are described by

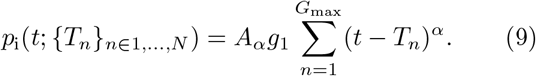

Here, *G*_max_ is the total number of infections. If *α* = 1, we have that

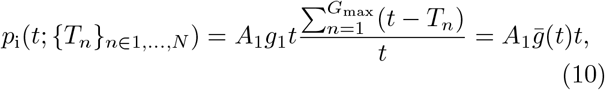

where 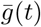 is the time-averaged phage genome concentration until time *t*. Hence, as noted by [11], when the protein concentration increases linearly in time, the protein accumulation is governed by the time-averaged genome concentration. Hence, the mean first passage time after multiple infections can be approximated by

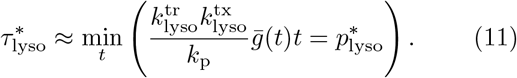

Note that infections that happen at times 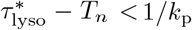 give rise to second order corrections. In the case of two infections, we find that 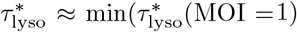, 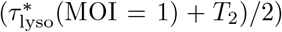, where 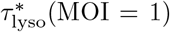 is the mean first passage time at an MOI of 1. Once 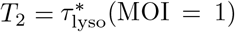, a decision has been reached, and the second infection does not affect the average outcome.

When the protein concentration increases with a temporal exponent *α >* 1, the mean no longer follows a simple scaling with 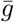. Phage genomes injected at earlier times contribute more to protein accumulation (Eq. (9)).Provided 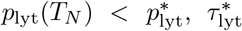 can be found using a quadratic equation (Appendix L). For two infections, we have

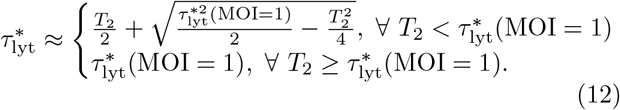

At small 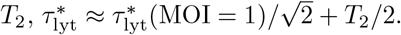.

Plotting 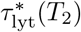 and 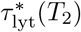 on the same axes (Fig. 2e), we can distinguish several regimes for the probability of lysogeny (Fig. 2f). As expected, at *T*_2_ = 0, the two phages infect simultaneously, and the preferred outcome is lysogeny 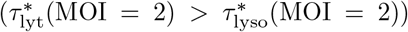. Initially, both curves in Fig. 2e increase with the same slope: 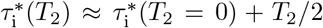. The curves hence stay parallel, and the decision is unchanged. However, as 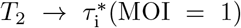, the curves plateau to levels 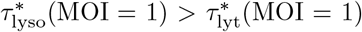. At the crossing point of the two 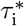 curves, the preferred outcome switches from lysogeny to lysis. After 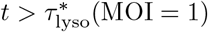, the decision is indistinguishable from a single infection.

## III. DISCUSSION

A temperate phage’s choice between lysis and lysogeny is an informed decision carried out by genetic systems of remarkable simplicity. We propose that the essential interactions underpinning these decision mechanisms can be identified from the temporal scaling behavior of their key components. In this framework, lysis-lysogeny decision mechanisms can be characterized by just a few parameters: the relative protein accumulation accelerations 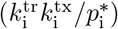; the timescale of the symmetry-breaking mechanism (e.g. 1*/k*_p_); as well as the temporal exponent of protein accumulation after symmetry-breaking (1 for *p*_lyso_). This approach illustrates how MOI-dependent decisions can arise despite all early gene transcription rates scaling in the same way with MOI [2, 12].

A key aspect of our model is that since temperate phages commit to either pathway within 5 min following infection, only processes that occur at rates faster than this timescale can influence the decision. We considered protease-mediated degradation of the lysogeny-inducing protein as an example, but our model also points toward alternative mechanisms. For example, enhanced degradation of the lysogeny-promoting mRNA species would cause *m* _lyso_ to reach a steady state soon after infection, and consequently, *p*_lyso_ would again increase linearly with time. Alternatively, if post-translational modification, e.g. phosphorylation, was required to activate the lytic protein, the concentration of activated lytic protein would increase with *t*^3^, again allowing for a crossing of the protein concentration curves.

In general, phages may have evolved to use multiple of these “building blocks” as the basis for their MOI-based decision-making. In phage *λ*, in addition to FtsH-mediated degradation of CII, RNase III destabilizes *cII* transcripts [3]. Nonlinearities in transcription, translation, or degradation rates could also affect decisions. For instance, although our model shows that saturation of host protease activity is not necessary for MOI-dependent decisions, it could nevertheless help stabilize lysogeny-promoting proteins (see Appendix M). This would be consistent with the *λ* protein CIII stabilizing CII by also being targeted by FtsH [27].

We find that our simple model neglecting all phage protein interactions at early infection stages also captures many stochastic features of phage decision making. At early times, protein concentration distributions appear Gaussian, meaning that first passage time distributions can be well approximated given the first and second moments. This approach is easily generalized for alternate symmetry-breaking mechanisms; master equations can be solved in the presence of fast mRNA degradation, protein activation, and other processes [28]. The first passage time distribution for proteins can then either be found exactly or using a clumping approximation. Finally, our model predicts a plateau in the probability of lysogeny: at high MOIs symmetry-breaking mechanisms become irrelevant, and first passage time variations are dominated by protein count noise.

Similarly, our scaling arguments for the effects of infection timings hold generally. By identifying the temporal exponents of protein accumulation, we can understand how mean first passage times scale with the timings of further infections. The probability of lysogeny follows from these temporal scalings (Fig. 2e,f). Intuitively, we find that once the second infection occurs after the mean first passage time at MOI = 1, the second infection no longer affects the outcome. This is consistent with earlier work in *λ* [12] suggesting that the second infection must occur in the activity window of CII.

Beyond MOI, phages can also base their decisions on host state. For example, a stress-induced transcription factor could directly couple either pathway to host conditions, without introducing an MOI-dependence. More subtly, an MOI-dependent mechanism, such as coupling to a host protease, could also inform phages of host state (Eq. (4)). Responding to new conditions including starvation, heat/osmotic/membrane stress, as well as quorum-sensing signals requires bacteria to extensively restructure their proteomes, thus reducing protease availability; phages can hence use proteases as a general indicator of host state, favoring lysogeny under stressful conditions [29–31]. Future experiments tracking protease activity together with lysogenization rates could quantitatively test our scaling predictions.

Our model suggests that the number of possible mechanisms for lysis-lysogeny decisions is quite limited, especially if coupling to essential host machinery is favored. This could explain why multiple phages base their decisions on the activity of FtsH [4, 6, 7], the only essential protease in *E. coli* [20, 32, 33]. Our framework provides concrete criteria for recognizing and classifying lysis-lysogeny decision mechanisms for both well-studied and novel temperate phages. Furthermore, our results suggest certain universal features of lysogenization probability curves, as well as the effects of infection timings.

## ACKNOWLEDGEMENTS

We thank Owen Abad, Ido Golding, Thu Vu Phuc Nguyen, and Martin Wühr for helpful discussions. This work was supported in part by Princeton University through the Center for the Physics of Biological Function, by grant NSF PHY-2309135 and the Gordon and Betty Moore Foundation Grant No. 2919.02 to the Kavli Institute for Theoretical Physics (KITP), and by the National Institute of General Medical Sciences (NIGMS) of the National Institutes of Health under award number 1R35GM122575. The content is solely the responsibility of the authors and does not necessarily represent the official views of the National Institutes of Health.

## CODE AVAILABILITY

The code for Gillespie simulations and figure generation can be found at github.com/harjuj/lysis_lysogeny_decisions.

## AUTHOR DECLARATIONS

The authors declare no competing interests.

## APPENDICES A MEAN DYNAMICS WITH mRNA AND PROTEIN DEGRADATION

For completeness, we may also include an mRNA degradation and dilution term 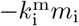 in Eq. (1). The equation can be solved [23] with the initial condition that *m*_i_(*t* = 0) = 0:

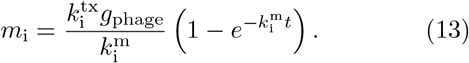

Similarly, we may include a protein degradation term 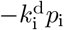 in Eq. (2). The solution is

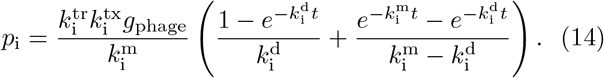

However, the leading order Taylor expansions for 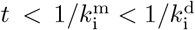, which are accurate at short times, recover the linear and quadratic dependence for mRNA and protein concentrations.

## B DISTANCE FROM THRESHOLD FOR REPRESSED PATHWAY

Suppose *g*_1_ and *g*_2_ correspond to phage genome concentrations at MOIs of 1 and 2, respectively; *g*_2_ = 2*g*_1_. Here, we derive an expression for how far the rejected pathway’s protein concentration is from its decision threshold when the other protein is at threshold levels. Let 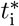 be defined by 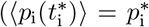. For simplicity, we assume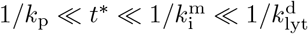.

First, at an MOI of 1, we have

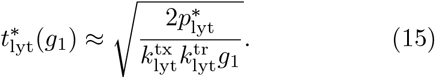

This implies that

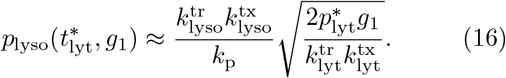

Similarly, at an MOI of 2, we have

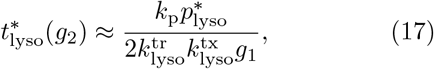

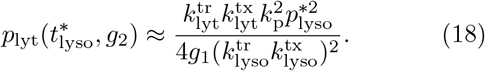

Equations (16) and (18) can be combined to find the relation

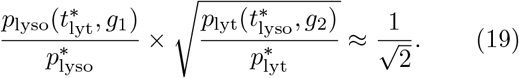

Eq. (19) describes a trade-off: if the concentration of lysogenic protein is low when the lytic decision is reached at an MOI of 1, the concentration of lytic protein will be close to the threshold when the lysogenic decision is reached at an MOI of 2. If we require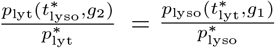, this ratio is given by 2^*−*1*/*3^ *≈* 0.8.

## C SCALING OF *t*_x_ WITH *k*_p_

The phage genome concentration *g*_lyso_ at which the expected lysis-lysogeny decision switches can generally be found by solving

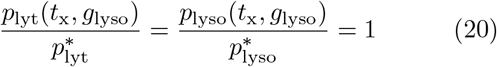

for *t*_x_ and *g*_lyso_. Here, we show that *t*_x_ = *t*^*\**^(*g*_lyso_) *∼* 1*/k*_p_, where *k*_p_ is the rate of protease degradation of the lysogeny protein. The scaling 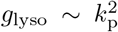 can then be derived as discussed in the main text.

Eq. (14) shows that even when mRNA and protein degradation and dilution are considered, *p*_i_ only depends linearly on *g*_phage_. This implies that the time at which the two curves cross, *t*_x_, is independent of *g*_phage_:

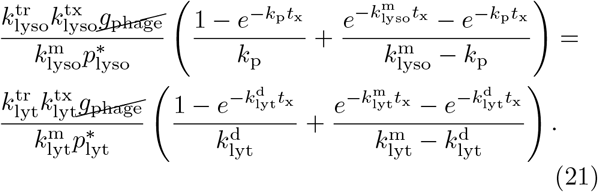

Hence, if a solution exists, *t*_x_ only depends on the rates *k*_p_, 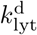, and 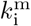, as well as on the ratio

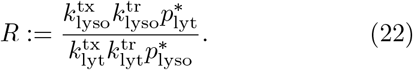

Graphically, as the MOI increases, both *p*_lyt_ and *p*_lyso_ move upwards on a log-log plot by the same amount; hence, their crossing point moves upwards along a straight line (Fig. 1c).

By defining 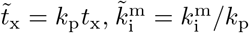 and 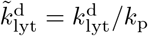 Eq. (21) can be written in the nondimensionalized form 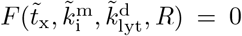. When *k*_p_ is significantly larger than 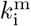 and 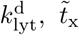, is approximated by the solution of 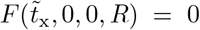. Consequently, *t*_x_ ∝ 1*/k*_p_, with a prefactor that only depends on *R*. This approximation breaks down as 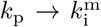 and the crossing time *t*_x_ approaches 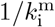. For decision mechanisms acting on timescales below 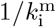, lysogeny would then be preferred (Fig. 1b).

## D PREFACTOR FOR Eq. (4) WHEN 1*/k*_p_ *« t*_x_

In the limit when 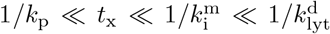, the prefactor for Eq. (4) can be found analytically. We write

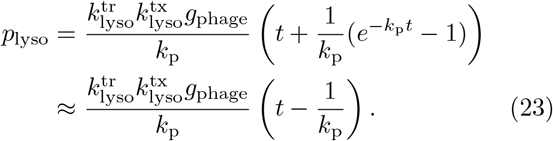

The curves cross when

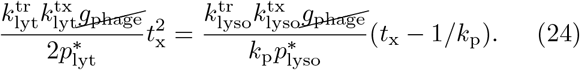

Neglecting the term 1*/k*_p_ gives a simple linear equation for *t*_x_ in the limit *t*_x_ *»*1*/k*_p_. Here, we include the linear correction by solving the quadratic equation. Dividing both sides by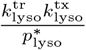, we recognize the ratio *R* (Eq. (22)), and solve for *t*_x_:

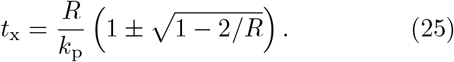

Note that we require *R >* 2 for a solution to exist; graphically, this corresponds to a requirement that the *p*_lyso_ curve is initially above the *p*_lyt_ curve (Fig. 1c). Since we expect a larger *R* to correspond to a larger initial distance between the *p*_i_ curves, *t*_x_ should increase with *R*. We hence take the positive branch of Eq. (25). The negative branch gives *t*_x_ ∼1*/k*_p_, breaking our assumption.

Substituting into 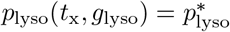 then gives

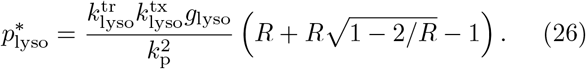

Solving for *g*_lyso_:

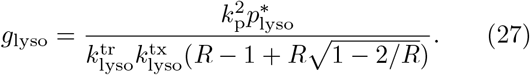

For 1*/k*_p_ ≲*t*_x_, *p*_lyso_(*t*) shows a transition from a linear to a quadratic temporal scaling (Fig. 1c), and the prefactor changes.

## E GILLESPIE SIMULATIONS

To study stochastic effects, we use Gillespie simulations to simulate trajectories of the number of mRNA copies *M*_i_ and the number of proteins *N*_i_. Let *G* be the MOI. The reactions we model are:

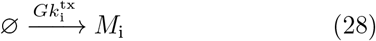

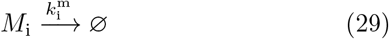

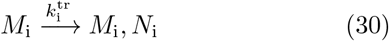

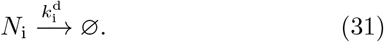

Note that the dynamics of the two protein species are independent. We further assume that the protein degradation rates are given by 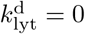 and 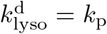. Results are similar for 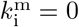 and 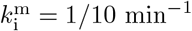, given that decisions are made within the first *∼*5 minutes.

For the population-average model, translation rates could be defined relative to the unknown threshold concentration levels. In stochastic simulations, the threshold concentrations are required, since we simulate absolute protein counts. Based on previous modeling work [19, 34], we use 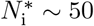.

Simulations were conducted using a custom Julia script.

## F TIME-DEPENDENT (*m*_lyt_, *p*_lyt_) DISTRIBUTION

First, we find the time-dependent probability distribution *P* (*M*_lyt_(*t*), *N*_lyt_(*t*)). For brevity, we omit the subscript lyt. At constant MOI *G*, the master equations for the system are:

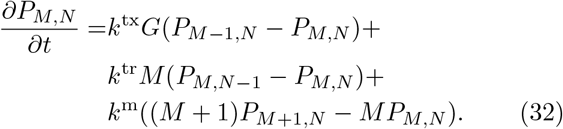

The steady-state distribution corresponding to this equation has been studied previously [22, 23]. Here, we are interested in the time-dependent dynamics of the distribution at times *t ≲*1*/k*^m^.

We define the generating function

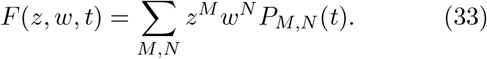

This gives rise to

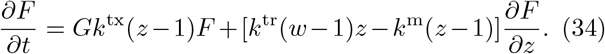

We define 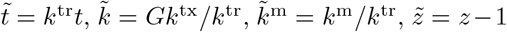and 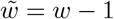. Dividing by *k*^tr^ and then by 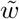 :

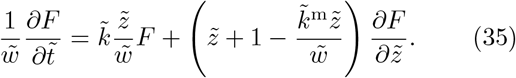

We use the method of characteristics to find a solution. 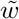 is a constant 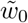 along the characteristic curve. We solve for 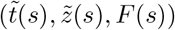 such that

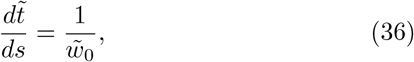

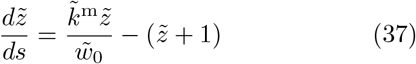

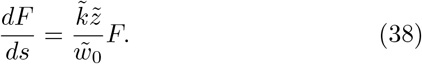

Integrating the first two equations, we find

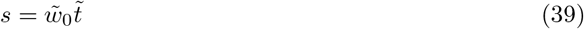

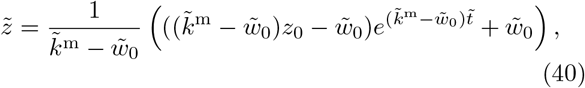

where 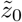is a constant value at *t* = *s* = 0. Note that we arbitrarily fix the starting point of the curve at 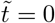.

*F* can be found by substituting Eq. (40) into Eq. (38):

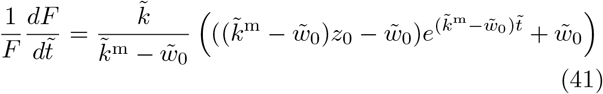

Integrating, we find

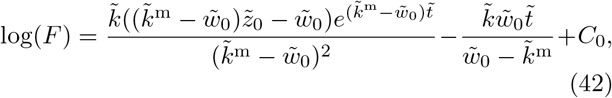

where *C*_0_ is a constant along each curve. By requiring that *F* (*t* = 0) = 1, we find

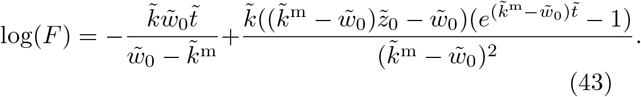

Substituting for 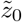 from Eq. (40), we have

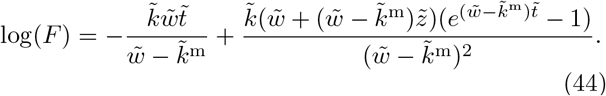

To simplify the expression, we substitute 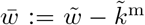. We then have

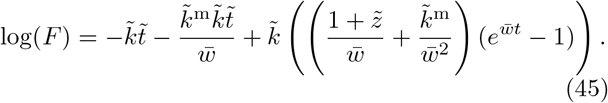

Up to a change of variables, this is the factorial cumulant generating function for the process; derivatives with respect to *z* and *t* yield factorial cumulants of *M* and *N* correspondingly. For mRNA, we find

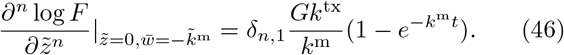

As expected, mRNA levels increase according to a Poisson process with rate *k*^tx^ while *t* «1*/k*^m^. For *t* » 1*/k*_m_, the distribution becomes a stationary Poisson distribution with rate *Gk*^tx^*/k*^m^.

To evaluate protein factorial cumulants, we expand Eq. (45) as a power series.

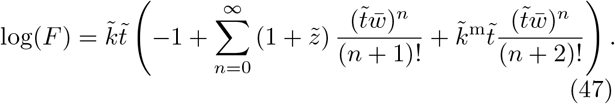

This form separates out *k*^m^ dependent terms; when *k*^m^ = 0, the second term in the summand is zero, and 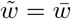. In this limit, protein factorial cumulants are given by

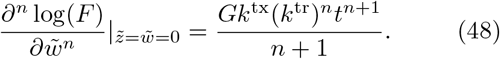

The mean and variance are

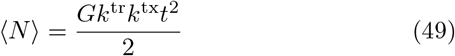

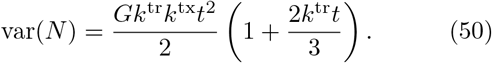

When 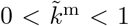, corrections to arbitrary order in 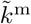can be found by only considering the relevant terms in Eq. (47). To first order we find:

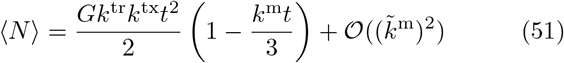

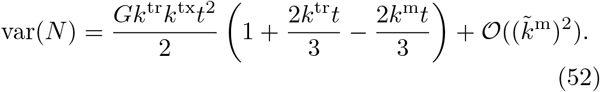

In practice, on timescales ∼ 5 min, where *k*^m^*t* ≈0.5, protein distributions are well-described by the zeroth order approximation.

## G TIME-DEPENDENT (*m*_lyso_, *p*_lyso_) DISTRIBUTION

Suppose we now introduce protein degradation into our model. We include a protease degradation term with rate *k*_p_, modeling the dynamics of quickly degraded *p*_lyso_. Again, we suppress the lyso subscript for brevity. Earlier, we saw that on relevant timescales, mRNA degradation introduces relatively small corrections to the protein distributions. As a simplification, we hence set *k*^m^ = 0. The generating function (Eq. (33)) now satisfies

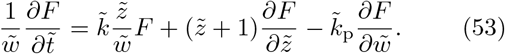

Here, 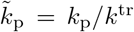. Note that 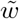 is no longer constant along the characteristic curve: we have

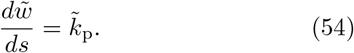

We find

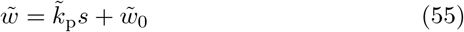

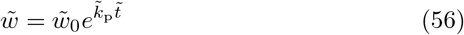

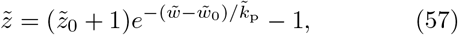

where 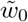 and 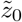 are constants for each curve, and we again set *t*(*s* = 0) = 0. log(*F* ) must now satisfy

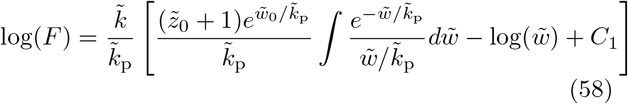

where *C*_1_ is a constant for each curve. This can be written in terms of the exponential integral Ei as:

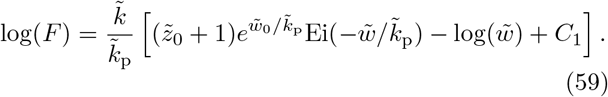

*C*_1_ can again be found by requiring that log(*F* (*t* = 0)) = 0. This yields

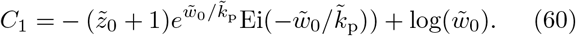

Our expression now contains the difference between two exponential functions:

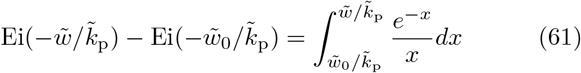

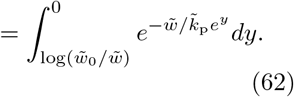

)

We then substitute for 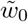 and 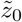:

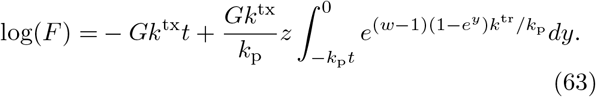

In the absence of mRNA degradation, *M* simply evolves as a Poisson process. For protein levels, we find

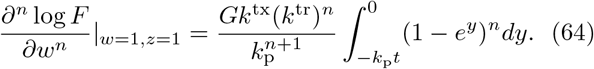

The mean and variance can now be found as

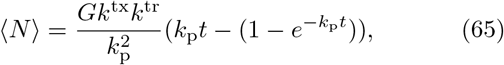

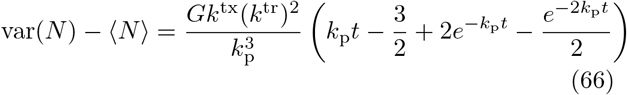

In the limit of *k*_p_*t →* 0, we recover our quadratic approximation. When *t »* 1*/k*_p_, *N* scales linearly with time.

## H PDF FOR 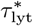

The 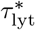 CDF (Eq. (6)) can be differentiated to find the PDF in the Gaussian regime. We define *z*(*t*) = (*p*^*\**^*−µ*(*t*))*/σ*(*t*), and write

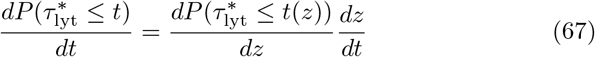

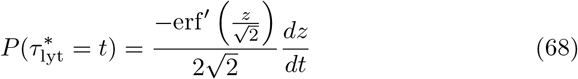

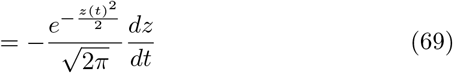

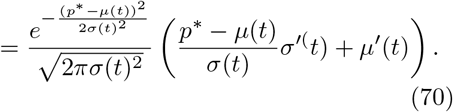

Here, we have suppressed the subscripts lyt for brevity. The mean *µ*(*t*) and standard deviation *σ*(*t*) are defined by Eq. (48) and (52). When mRNA degradation is negligible, the derivatives can be found as:

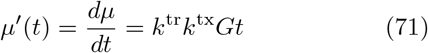

and

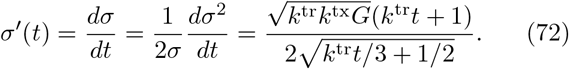

## I EXPECTATION VALUE FOR 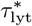

The PDF is dominated by the Gaussian peak around time 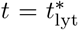 where the exponent *z* is zero. We can find the mean first passage time as

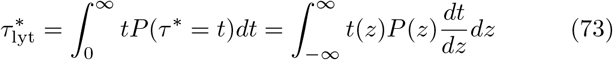

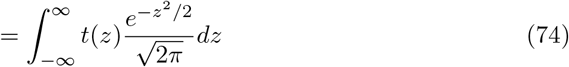

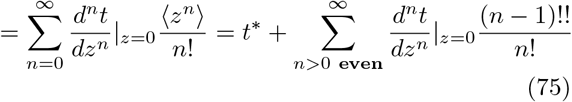

where we used Eq. (69). *z ∼ N* (0, 1), which implies its even moments are given by *σ*^*p*^(*p −* 1)!!, where !! denotes . the double factorial.

To leading order, we have

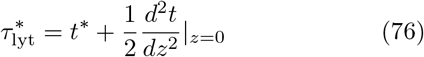

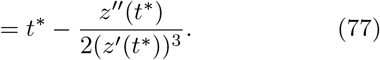

We have

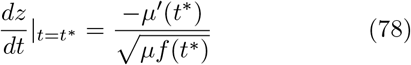

where *f* (*t*) = (1 + 2*k*^tr^*t/*3) is the Fano factor. Using *µ* (*t*) = 2*µ/t*, we write

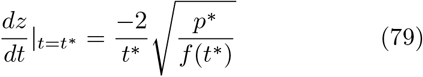

and

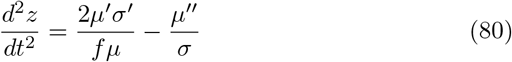

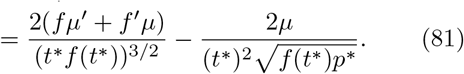

We hence have

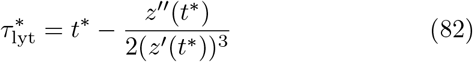

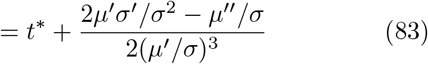

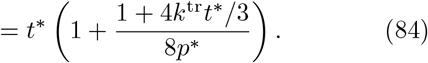

The correction can be written as

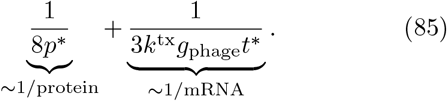

Hence, when protein and mRNA counts are of order ∼ 10 at the time the threshold is reached, both corrections are small, and *τ*^*\**^≈*t*^*\**^. For small *g*, the increase in the mean first passage time is due to fluctuations in mRNA counts. For large *g*_phage_, the noise is simply due to the fluctuations in protein counts near the threshold; var (*N*) ∼ *N* . An increase in *g* no longer affects the noise level around the threshold.

We hence predict that for

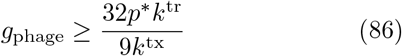

counting noise dominates, and 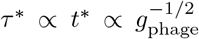 . For small *g*_phage_, there is a correction that scales as 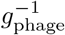.

## J MOI-DEPENDENCE DUE TO NOISE

Suppose that neither protein is degraded by a protease, and that near 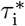, protein concentrations are well-approximated by a Gaussian. We further assume that *τ*_lyso_ *< τ*_lyt_ (this ensures that the mean curves cross when *k*_p_ *>* 0, as shown in Fig. 1c).

The probability for lysogeny can be found by substituting Eq. (6) and (70) into Eq. (5). As shown in Fig. 2D, of *k* this prediction agrees with simulations, and in the case of 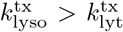, predicts a slight decrease in ℙ_lyso_ at low MOIs. Here, we provide an intuitive explanation for why this dependence emerges.

Consider

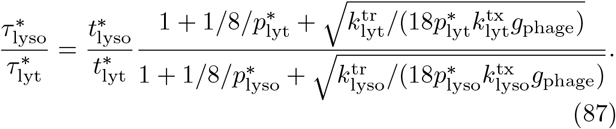

When

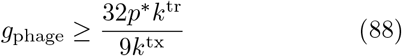

the fraction is independent of the MOI:

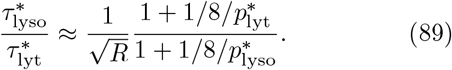

Hence, at large MOIs, we expect the probability of lysogeny to plateau, as the protein count noise near the threshold approaches a constant for both proteins.

At small *g*_phage_, we have

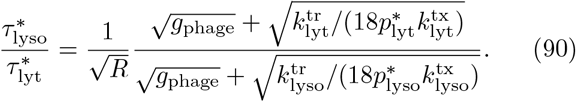

Let 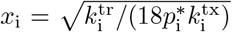. As *g*_phage_ decreases, the fraction increases if *x*_lyt_ *> x*_lyso_. Consider three possibilities consistent with 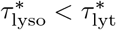:

1. 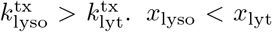, so the fraction increases as *g*_phage_ decreases, and lysis events become more likely, as shown in Fig. 2d.
2. 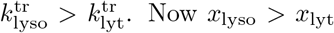. Now *x*_lyso_ *> x*_lyt_, so the fraction decreases. Intuitively, a smaller mRNA count is required for *p*_lyso_ to reach the threshold, so mRNA count noise at low MOIs does not slow down the accumulation of *p*_lyso_ as much.
3. 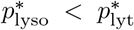. Again, the fraction decreases, and lysogeny can become more favored at low MOIs. Similar to before, a smaller mRNA count is required to reach the threshold, and hence mRNA count noise is not as relevant. Note, however, that protein count noise increases 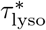 more at large MOIs; we hence expect a lower rate of lysogeny at large MOIs.

## K CALCULATING 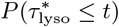

The concentration curves *p*_lyso_ typically increase quickly with time, and show occasional downwards steps due to protein degradation. The first passage time problem can hence be simplified by assuming that: (1) crossings of 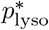 are “clumped”, i.e. there is a typical number of crossings that happen in relatively quick succession; and (2) the protein level will not drop significantly at later points in time, so that each trajectory only contains one such clumped crossing (Supplemental Figure S3 a,b). The number of clumped crossings is hence a binary variable (0 or 1), and the probability of a clumped crossing in a given time-interval is given by the mean:

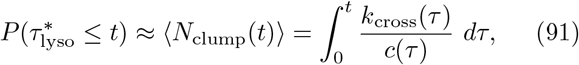

where *N*_clump_(*t*) ∈{0, 1}is the number of clumps within the time *t*.

To calculate *k*_cross_(*t*) (Eq. (7)), the mean rate at which *p*_lyso_ levels cross the threshold 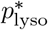, we require the conditional mean 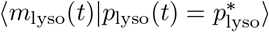 . In the Gaussian regime, both protein and mRNA levels are jointly normally distributed. Conditional probabilities can then be calculated given the second moments of the distribution.

We use Eq. (63) to calculate

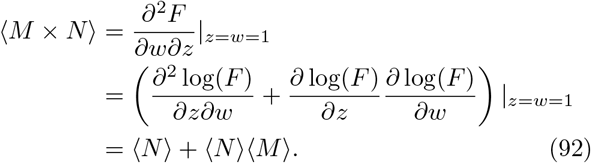

We then have

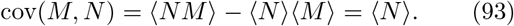

Combining this with the variance defined by Eq. (66), the conditional expectation is then given by:

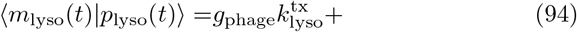

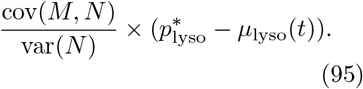

Next, we derive the clump size *c*(*t*); the number of expected crossings when 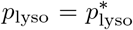. Let 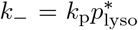 be the degradation probability at 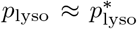, and let 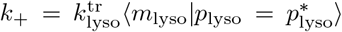 be the probability of a translation event.

First, consider *k*_+_ *> k*_*−*_ . Consecutive crossings are expected to occur every time the protein count falls down by one from the threshold before climbing up (Supplemental Figure S3c). This approximation neglects cases where the protein level climbs further up before falling below the threshold.

The probability of falling down below the level before a climbing step is *p* = *k*_*−*_ */k*_+_. The probability of escaping *p*^*\**^ is given by (1*− p*_*−*_ ). The expected number of crossings is hence given by the expected number of trials before a successful escape, namely, 1*/*(1*− p*_*−*_ ) by the geometric distribution.

By contrast, if *k*_+_ *< k*_*−*_, a crossing can occur again from the downward direction with probability *k*_+_*/k*_*−*_ . Similar to before, the number of trials before the protein count escapes to levels below *p*^*\**^ is then given by 1*/*(1 *− p*_+_). Combining the two expressions gives

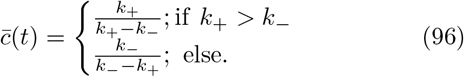

Combining these two expressions and substituting the rates of translation and protein degradation gives:

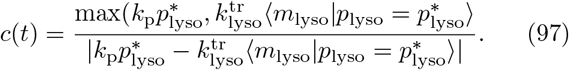

## L 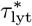 WITH MULTIPLE INFECTIONS

Given multiple infections at times {*T*_*n*_} _*n* ∈1,…,*N*_ the mean concentration *p*_lyt_ at time *T*_*N*_ *< t* is given by

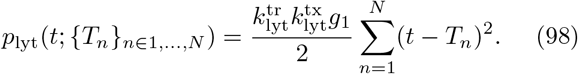

By grouping terms by order of *t*, we find

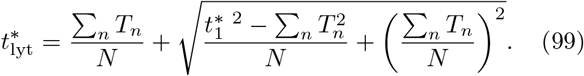

As discussed in Section I, 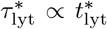. Multiplying both sides by the prefactor 1 + *f*, and by defining 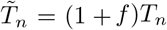, we find

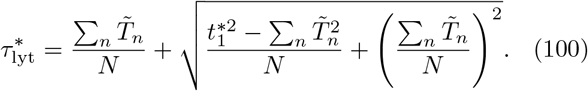

As *T*_*N*_ →0, we have that 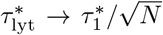, as expected. When *f* is small, setting *N* = 2 recovers Eq. (12). The difference between *τ* ^*\**^ and *t*^*\**^ results in infections giving contributions that appear “delayed” by a factor (1 + *f* ), similar to how the expectation value *τ* ^*\**^ is delayed by the same factor from the deterministic expectation *t*^*\**^.

## M SATURATION OF PROTEASE ACTIVITY

Suppose protease-mediated degradation is initially linear (Eq. (3)) but saturates to a constant level once *p*_lyso_(*t*_sat_) = *p*_sat_, where *t*_sat_ is the saturation time and *p*_sat_ a saturation threshold concentration. The concentration curve *p*_lyso_(*t*) can now have three scaling regimes: (1) *p*_lyso_ ∼*t*^2^ when *t <* 1*/k*_p_; (2) *p*_lyso_ ∼*t* when 1*/k*_p_ *< t < t*_sat_; and (3) *p*_lyso_ ∼ *t*^2^ once protease activity saturates for *t > t*_sat_. At higher MOIs, *p*_sat_ is reached faster, and the duration of regime (2) decreases.

If 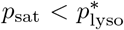, the lytic pathway is destabilized: once *p*_lyso_(1*/k*_p_) *> p*_sat_, regime II ceases to exist, and lysogeny is preferred (Fig. 1b). Furthermore, if a lysis decision is reached at time *τ* ^*\**^, *p*_lyso_(*τ* ^*\**^) is higher than in the absence of protease activity saturation.

If 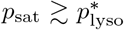, the lysogenic pathway is stabilized. In this case, protease activity saturation does not significantly affect *p*_i_(*t*) for *t < τ* ^*\**^. Hence, the same MOI-based decision is reached. If the lytic decision is reached and *p*_lyso_ is degraded before reaching *p*_sat_, the lytic pathway remains stable. By contrast, if a lysogeny decision is made, *p*_lyso_ reaches saturation levels, enhancing its net accumulation after the decision.

## References

[1] P. Kourilsky, Lysogenization by bacteriophage lambda, Molecular and General Genetics MGG 122, 183 (1973).

[2] A. B. Oppenheim, O. Kobiler, J. Stavans, D. L. Court, and S. Adhya, Switches in Bacteriophage Lambda Development, Annual Review of Genetics 39, 409 (2005).

[3] S. R. Casjens and R. W. Hendrix, Bacteriophage lambda: Early pioneer and still relevant, Virology 60th Anniversary Issue, 479-480, 310 (2015).

[4] C. Herman, T. Ogura, T. Tomoyasu, S. Hiraga, Y. Akiyama, K. Ito, R. Thomas, R. D’Ari, and P. Bouloc, Cell growth and lambda phage development controlled by the same essential Escherichia coli gene, ftsH/hflB., Proceedings of the National Academy of Sciences 90, 10861 (1993).

[5] Y. Shotland, S. Koby, D. Teff, N. Mansur, D. A. Oren, K. Tatematsu, T. Tomoyasu, M. Kessel, B. Bukau, T. Ogura, and A. B. Oppenheim, Proteolysis of the phage lambda CII regulatory protein by FtsH (HflB) of Escherichia coli, Molecular Microbiology 24, 1303 (1997).

[6] G. W. Broussard, L. M. Oldfield, V. M. Villanueva, B. L. Lunt, E. E. Shine, and G. F. Hatfull, Integration-Dependent Bacteriophage Immunity Provides Insights into the Evolution of Genetic Switches, Molecular Cell 49, 237 (2013).

[7] I. M. Murchland, A. Ahlgren-Berg, J. M. J. Pietsch, A. Isabel, I. Dodd, and K. E. Shearwin, Instability of CII is needed for efficient switching between lytic and lysogenic development in bacteriophage 186, Nucleic Acids Research 48, 12030 (2020).

[8] J. S. Weitz, Y. Mileyko, R. I. Joh, and E. O. Voit, Collective Decision Making in Bacterial Viruses, Biophysical Journal 95, 2673 (2008).

[9] M. Avlund, I. B. Dodd, K. Sneppen, and S. Krishna, Minimal Gene Regulatory Circuits that Can Count like Bacteriophage Lambda, Journal of Molecular Biology 394, 681 (2009).

[10] M. L. Robb and V. Shahrezaei, Stochastic Cellular Fate Decision Making by Multiple Infecting Lambda Phage, PLOS ONE 9, e103636 (2014).

[11] M. G. Cortes, J. T. Trinh, L. Zeng, and G. Balázsi, Late-Arriving Signals Contribute Less to Cell-Fate Decisions, Biophysical Journal 113, 2110 (2017).

[12] T. Yao, S. Coleman, T. V. P. Nguyen, I. Golding, and O. A. Igoshin, Bacteriophage self-counting in the presence of viral replication, Proceedings of the National Academy of Sciences 118, e2104163118 (2021).

[13] M. Lieb, The establishment of lysogeny in Escherichia coli, Journal of Bacteriology 65, 642 (1953).

[14] F. St-Pierre and D. Endy, Determination of cell fate selection during phage lambda infection, Proceedings of the National Academy of Sciences of the United States of America 105, 20705 (2008).

[15] L. Zeng, S. O. Skinner, C. Zong, J. Sippy, M. Feiss, and I. Golding, Decision making at a subcellular level determines the outcome of bacteriophage infection, Cell 141, 682 (2010).

[16] J. T. Trinh, T. Székely, Q. Shao, G. Balázsi, and L. Zeng, Cell fate decisions emerge as phages cooperate or compete inside their host, Nature Communications 8, 14341 (2017).

[17] I. Golding, Single-Cell Studies of Phage λ: Hidden Treasures Under Occam’s Rug, Annual Review of Virology 3, 453 (2016).

[18] Q. Shao, J. T. Trinh, and L. Zeng, High-resolution studies of lysislysogeny decision-making in bacteriophage lambda, Journal of Biological Chemistry 294, 3343 (2019).

[19] A. Arkin, J. Ross, and H. H. McAdams, Stochastic Kinetic Analysis of Developmental Pathway Bifurcation in Phage λ-Infected Escherichia coli Cells, Genetics 149, 1633 (1998).

[20] M. Gupta, A. N. T. Johnson, E. R. Cruz, E. J. Costa, R. L. Guest, S. H.-J. Li, E. M. Hart, T. Nguyen, M. Stadlmeier, B. P. Bratton, T. J. Silhavy, N. S. Wingreen, Z. Gitai, and M. Wühr, Global protein turnover quantification in Escherichia coli reveals cytoplasmic recycling under nitrogen limitation, Nature Communications 15, 5890 (2024).

[21] V. Shahrezaei and P. S. Swain, Analytical distributions for stochastic gene expression, Proceedings of the National Academy of Sciences 105, 17256 (2008).

[22] M. Thattai and A. van Oudenaarden, Intrinsic noise in gene regulatory networks, Proceedings of the National Academy of Sciences 98, 8614 (2001).

[23] P. Bokes, J. R. King, A. T. A. Wood, and M. Loose, Exact and approximate distributions of protein and mRNA levels in the low-copy regime of gene expression, Journal of Mathematical Biology 64, 829 (2012).

[24] K. R. Ghusinga, J. J. Dennehy, and A. Singh, First-passage time approach to controlling noise in the timing of intracellular events, Proceedings of the National Academy of Sciences 114, 693 (2017).

[25] K. Rijal, A. Prasad, A. Singh, and D. Das, Exact Distribution of Threshold Crossing Times for Protein Concentrations: Implication for Biological Timekeeping, Physical Review Letters 128, 048101 (2022).

[26] D. Aldous, in Probability Approximations via the Poisson Clumping Heuristic, edited by D. Aldous (Springer, New York, NY, 1989) pp. 1–22, 87–89.

[27] C. Herman, D. Thévenet, R. D’Ari, and P. Bouloc, The HflB protease of Escherichia coli degrades its inhibitor lambda cIII., Journal of Bacteriology 179, 358 (1997).

[28] T. Jahnke and W. Huisinga, Solving the chemical master equation for monomolecular reaction systems analytically, Journal of Mathematical Biology 54, 1 (2007).

[29] M. Obuchowski, Y. Shotland, S. Koby, H. Giladi, M. Gabig, G. Wegrzyn, and A. B. Oppenheim, Stability of CII is a key element in the cold stress response of bacteriophage lambda infection, Journal of Bacteriology 179, 5987 (1997).

[30] M. Somiska, P. Neubauer, and G. Wgrzyn, Regulation of Bacteriophage λ Development by Guanosine 5-Diphosphate-3-diphosphate, Virology 262, 431 (1999).

[31] Y. Geng, T. V. P. Nguyen, E. Homaee, and I. Golding, Using bacterial population dynamics to count phages and their lysogens, Nature Communications 15, 7814 (2024).

[32] T. Ogura, K. Inoue, T. Tatsuta, T. Suzaki, K. Karata, K. Young, L.-H. Su, C. A. Fierke, J. E. Jackman, C. R. H. Raetz, J. Coleman, T. Tomoyasu, and H. Matsuzawa, Balanced biosynthesis of major membrane components through regulated degradation of the committed enzyme of lipid A biosynthesis by the AAA protease FtsH (HflB) in Escherichia coli, Molecular Microbiology 31, 833 (1999), _eprint: https://onlinelibrary.wi-ley.com/doi/pdf/10.1046/j.1365-2958.1999.01221.x.

[33] L.-M. Bittner, J. Arends, and F. Narberhaus, When, how and why? Regulated proteolysis by the essential FtsH protease in Escherichia coli, Biological Chemistry 398, 625 (2017).

[34] R. I. Joh and J. S. Weitz, To Lyse or Not to Lyse: Transient-Mediated Stochastic Fate Determination in Cells Infected by Bacteriophages, PLOS Computational Biology 7, e1002006 (2011).

