## Supplementary Figures for "Counting to two: how phages decide between lysis and lysogeny"

(Dated: July 24, 2026)

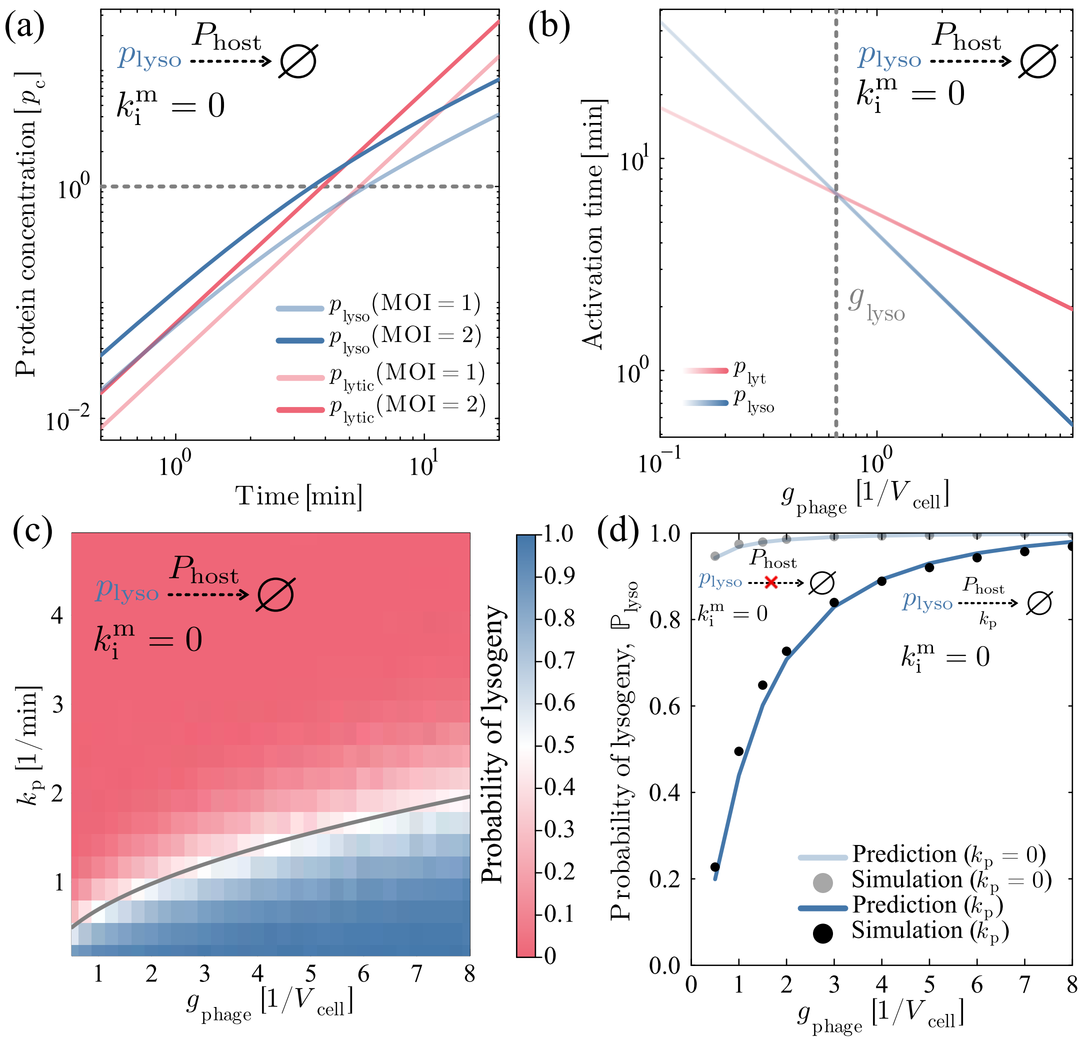

FIG. S1. **Model without mRNA degradation.** (a) Mean protein concentration curves in the absence of mRNA degradation, similar to Main Fig. 1c. (b) Estimated activation times in the absence of mRNA degradation, similar to Main Fig. 1d. (c) Probability of lysogeny with varying MOI and protease activity rate in Gillespie simulations without mRNA degradation, similar to Main Fig. 2b. (d) Probability of lysogeny together with analytical predictions as a function of MOI, similar to Main Fig. 2d.

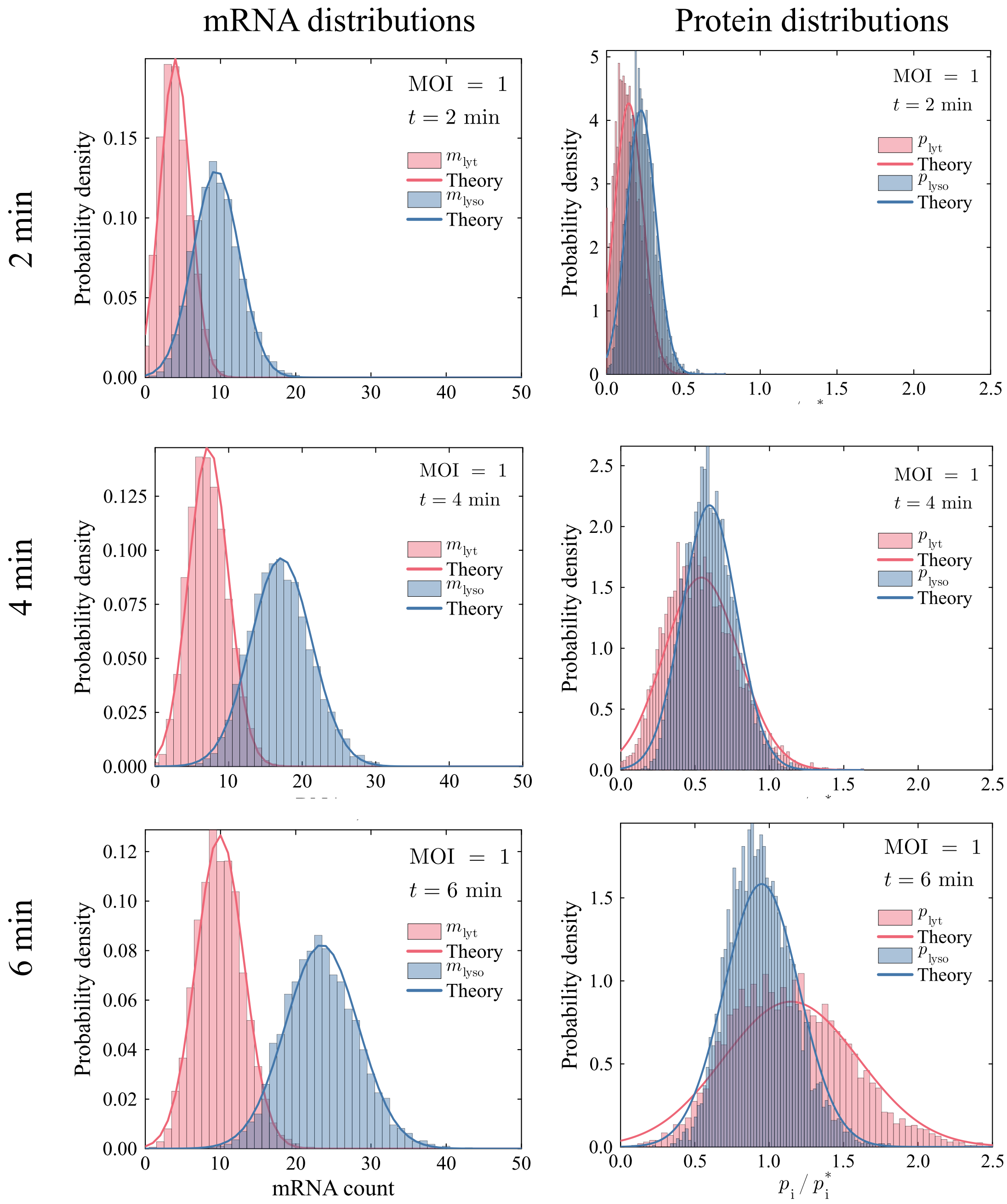

FIG. S2. **mRNA and protein distributions.** Table of histograms for mRNA (left) and protein (right) counts in Gillespie simulations. Rows show simulation data for $t = 2,\ 4,\ 6$ min post-infection. Overlaid curves show Gaussian curves with theoretically derived mean and standard deviation.

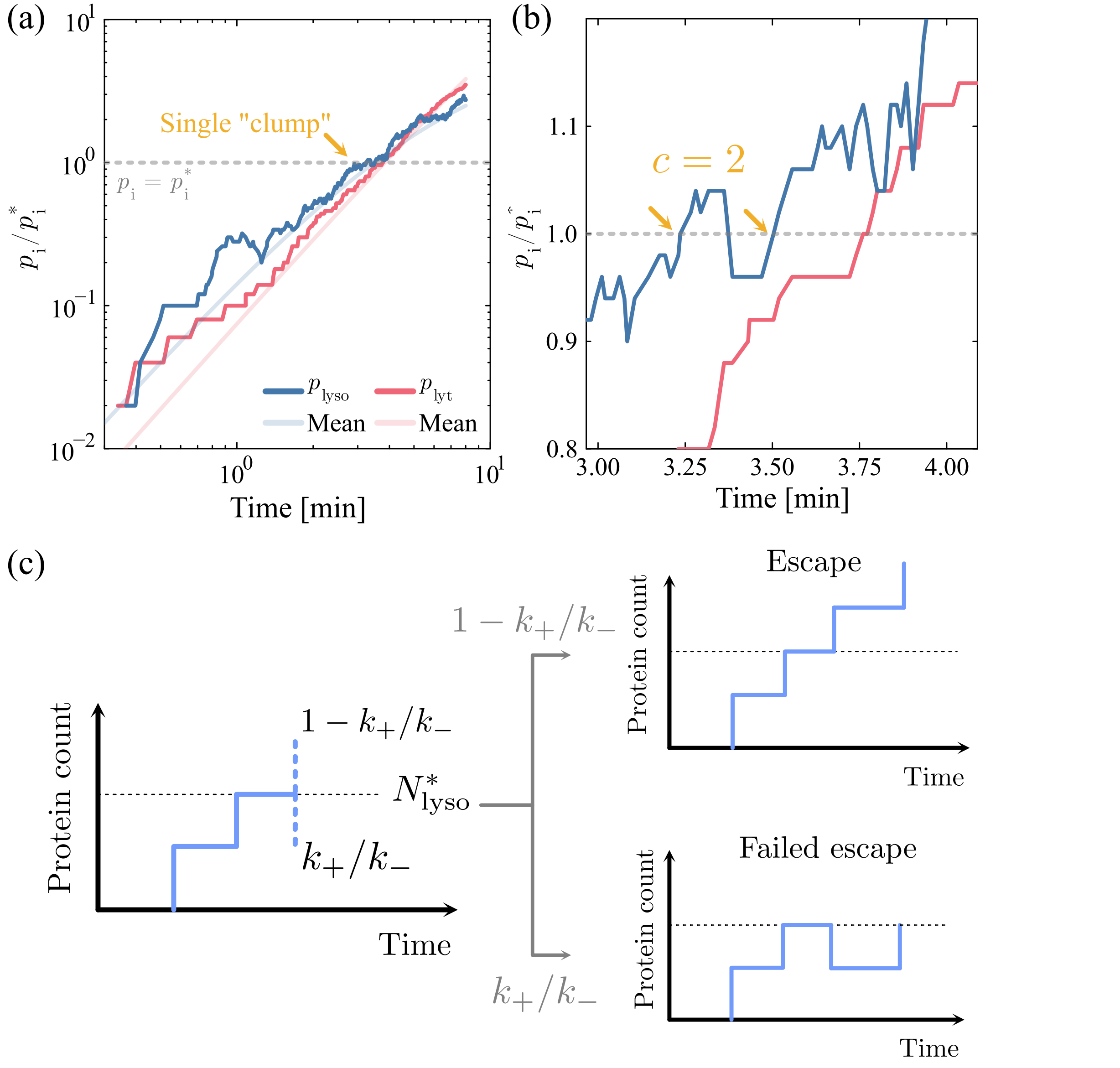

FIG. S3. **Clump rate approximation.** (a) Example protein concentration trajectory from Gillespie simulations at MOI = 2. Orange arrow indicates crossing point where $p_{lyso} \approx p^*_{lyso}$, corresponding to a clump of crossings that occur at similar times. Note that due to strong deterministic drift, we only expect to observe a single clump during a trajectory. (b) Zoomed in view of the trajectories in panel (a). Orange arrows show the two crossing points where $p_{lyso} = p^*_{lyso}$. In this particular case, the clump size is two upward crossings. (c) Illustration of the geometric distribution approximation. Once $p_{lyso}$ is at the critical level, there are two possibilities; either the protein concentration drops downwards (failed escape) in which case another crossing will occur. Alternatively, the protein level increases, which we consider a successful escape. Note that this approximation neglects cases where the protein count has increased by two and then drops below the decision threshold again.
